# Distinctive diffusive properties of swimming planktonic copepods in different environmental conditions

**DOI:** 10.1101/343582

**Authors:** Raffaele Pastore, Marco Uttieri, Giuseppe Bianco, Maurizio Ribera d’Alcalá, Maria Grazia Mazzocchi

## Abstract

Suspensions of small planktonic copepods represent a special category in the realm of active matter, as their size falls within the range of colloids, while their motion is so complex that it cannot be rationalized according to basic self-propelled particle models. Indeed, the wide range of individual variability and swimming patterns resemble the behaviour of much larger animals. By analysing hundreds of three dimensional trajectories of the planktonic copepod *Clausocalanus furcatus* we investigate the possibility of detecting how the motion of this species is affected by different external conditions, such as the presence of food and the effect of gravity. While this goal is hardly achievable by direct inspection of single organism trajectories, we show that this is possible by focussing on simple average metrics commonly used to characterize colloidal suspensions, such as the mean square displacement and the dynamic correlation functions. We find that the presence of food leads to the onset of a clear localization that separates a short-time ballistic from a long-time diffusive regime. Such a benchmark reflects the tendency of *C. furcatus* to remain temporally feeding in a limited space and disappears when food is absent. Localization is clearly evident in the horizontal plane, but is negligible in the vertical direction, due to the effect of gravity. Our results suggest that simple average descriptors may provide concise and useful information on the swimming properties of planktonic copepods, even though single organism behaviours are strongly heterogeneous.

## 1 Introduction

Motion behaviour reveals the dynamic adaptations of an organism to a changing environment. Movement can have direct consequences on the fitness of the individual, but it can also affect up to the population and metapopulation levels ^1^^;^^2^. The first model used to simulate organism movement was the random walk (see, *e.g.* Ref.s^3^^;^^4^). However, this was demonstrated insufficient to represent biological systems that are intrinsically characterized by a high degree of complexity. As such, animal movement may display regular patterns that cannot be assimilated to random walks^5^^;^^6^. In some instances, the emergence of regularities depends on the considered timescale and might manifest an adaptive behaviour for organism’s survival. Indeed, movement is crucial for organism’s life and represents a tradeoff between searching for food and mates, and escaping from predators (^7^ and references therein). In water, movement is constrained by the medium since friction is higher as compared to the air in terrestrial environments. Therefore, water acts as a selective force favoring specific organismal morphologies *e.g.^8^* and reduces significantly animal’s speed. Notwithstanding this constraint, small planktonic metazoans, living in the water at the edge between viscous and inertial environments^9^, display a large suite of swimming behaviors^10^^;^^11^. The various motion patterns of planktonic copepods, the most numerous animals at sea, have been categorised in two prevalent modes, *i.e.* “ballistic” and “diffusive” ^12^. In many active systems, these two modes interplay on different time and length scales, and are often tangled with intermittent individual dynamics, so as to optimise target search ^13^^;^^14^^;^^15^^;^^16^. Intermittency in planktonic copepod trajectories has been, indeed, observed in recent experiments, which also unveiled the presence of regularities contributing to fitness and likely to the observed behavioural diversity^6^.

To shed light on the adaptive *vs.* neutral component of copepod motion behaviour, we carried out a quantitative analysis of the swimming trajectories of a prominent planktonic species, *Clausocalanus furcatus* which is commonly distributed in warm epipelagic waters and displays unique swimming patterns^6^. Our goal is to understand which common properties emerge from the heterogeneity of individual motions and whether these features are useful benchmarks to distinguish different external conditions, namely the presence or absence of food, and to discern the effect of gravity. To this aim, we exploited the quantitative tools and the ensemble averaged descriptors routinely used to characterize the properties of soft matter systems, such as simple liquids and colloidal suspensions. Monitoring the mean square displacement (MSD) and the dynamic correlation functions, we show that, in the presence of food, an intermediate time plateau appears, which separates a short-time ballistic regime from a longtime diffusion. This plateau emerges from localized motion in the horizontal plane and is not observed in the vertical plane where gravity acts. Moreover, localization does not appear when the copepods move in the absence of food particles. These results suggest that *C. furcatus* is able to finely tune their swimming patterns depending on the external conditions, and that the presence of temporary localized motion is a good benchmark to distinguish among them. In the final part of the paper, we discuss the emerging scenario in an ecological context.

## 2 Methods

### 2.1 Clausocalanus furcatus

*Clausocalanus furcatus* is a calanoid copepod with total body length ≃ 1 *mm* and is widespread in epipelagic waters of both oligotrophic^17^^;^^18^^;^^19^ and coastal eutrophic regions ^20^^;^^21^^;^^22^. A unique feature of this species is its incessant swimming behaviour, consisting of a sequence of high-speed movements ≃ 10 *mm* s^−1^) on short timescales, which assemble in more complex and convoluted patterns on longer time scale^6^^;^^10^^;^^23^. Typically, this species captures prey items falling within a narrow frontal capture area^23^, using its chemical and mechanical sensory structures^24^.

### 2.2 Experiments

Trajectories of *C. furcatus* adult females were recorded during the experiments described in details in^6^. Briefly, zooplankton samples were collected during the autumn of the years 2008 and 2009 from vertical tows with a 200 *µm* mesh net in the upper 50 *m* of the water column in the Gulf of Naples (Tyrrhenian Sea, Western Mediterranean). Soon after collection, samples were brought to the laboratory, where individuals of the target species were sorted from the samples and checked under a dissecting microscope for stage and sex. Healthy females were selected and separated in homogeneous groups (*n* = 30 − 37) for the experiments and always kept at the same temperature of sampling site (19 − 21° *C).* In each year, two recordings were performed lasting one hour each: Exp1 and Exp2 in 2008, Exp3 and Exp4 in 2009. Exp1 and Exp3 were recorded in the presence of food, which consisted of natural particle assemblages collected at the same time and site of zooplankton sampling. Food concentration was *ρ_f_* = 5 · 10^5^ cells *L*^−1^ and 5 · 10^6^ cells *L*^−1^ for Exp1 and Exp3, respectively. Exp2 and Exp4 were, instead, recorded in the absence of food (using filtered sea water). Three-dimensional (3D) trajectories were obtained using an automatic computer vision system, composed by two orthogonally positioned digital cameras, equipped with telecentric lenses and two backlight infrared lamps. The spatial resolution of the system was 78 *µm* and images were acquired at 15*fps.* To avoid any impact of visible light on the copepod behavior, recordings were performed with a red light illumination and in sufficiently large aquarium (1 *L*). The field of view of the system was set to be 10 *mm* away from the aquarium walls and 20 *mm* from its bottom and the water surface. Individuals exiting and entering successively in the field of view of the system were recorded as separate tracks. The number of trajectories recorded during each experiment was 255 for Exp1, 384 for Exp2, 625 for Exp3 and 425 for Exp4. Overall, recorded trajectories had minimum duration of 5 *s* average duration of 28 *s* and maximum duration of 5 *min* and 42 *s.*

### 2.3 Analysis of trajectories

To investigate the properties of *C. furcatus* swimming, we focused on simple average descriptors that are commonly used to characterize liquids, colloidal suspensions and other soft matter systems. These include the mean square displacement (MSD) in 3D, 〈Δ*r*^2^(Δ*t*)〉 = 〈Δ*x*^2^(Δ*t*) + Δ*y*^2^(Δ*t*) + Δ*z*^2^(Δ*t*)〉, as well as its components in the horizontal plane, 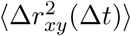, and in the vertical direction, 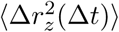. Averages were computed both over time and individual trajectories. See Sec.5 for further technical details. The MSD is the simplest and most intuitive metric to characterize the motion of a system from particle tracking data. It enables to estimate the average distance an organism moved from its position at an arbitrary time origin, after a time interval Δ*t*.

In addition, we investigated the dynamic correlation function relative to displacements in the horizontal plane, 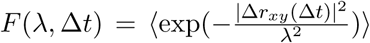 This function is akin to the intermediate self scattering function, as it estimates the average fraction of organisms that, after a lag-time Δ*t*, moved less than a probe length *λ*^25^. For any value of *λ* this function must be one in the short time limit, lim_Δ*t*→0_ *F*(*λ* Δ*t*) = 1, whereas it must vanish at long time, lim_Δ*t*→∞_ *F*(*λ* Δ*t*) = 0, as far as the investigated system is not arrested on the length scale of *λ.* Finally, we measured the relaxation time *τ_λ_* of *C. furcatus* from the decay of *F*(*λ* Δ*t*), by using the standard definition *F*(*λ, τ_λ_*) = *e*^−1^. In general, *τ_λ_* ∝ *λ* is expected for ballistic systems and *τ_λ_* ∝ *λ*^2^ for diffusive systems.

## 3 Results

Figure 1 illustrates some examples of single *Clausocalanus furcatus* trajectories from two experiments in the presence (panel a) and absence of food (panel b). Figure 2 a and b display instances of spatial coordinates recorded in the horizontal and in the vertical direction, as well as 3D square displacements as a function of the time-lag after the first recorded point in each trajectory, Δ*r_i_*(Δ*t*), respectively. These figures summarize the scenario emerging from previous studies^6^^;^^23^^;^^26^: a variety of moving patterns in *C. furcatus* with the possibility of recognizing some peculiar and recursive swimming behaviour. However, in lack of robust quantitative benchmarks and due to the large individual heterogeneity, establishing clear-cut relations between these behaviour and the different experimental conditions appears to be more difficult. Despite this, if any property of the motion becomes particularly relevant and emerges on top of the individual variability, we do expect that calculating average quantities on a large number of trajectories is the simplest way to highlight these features.

**Figure 1:**
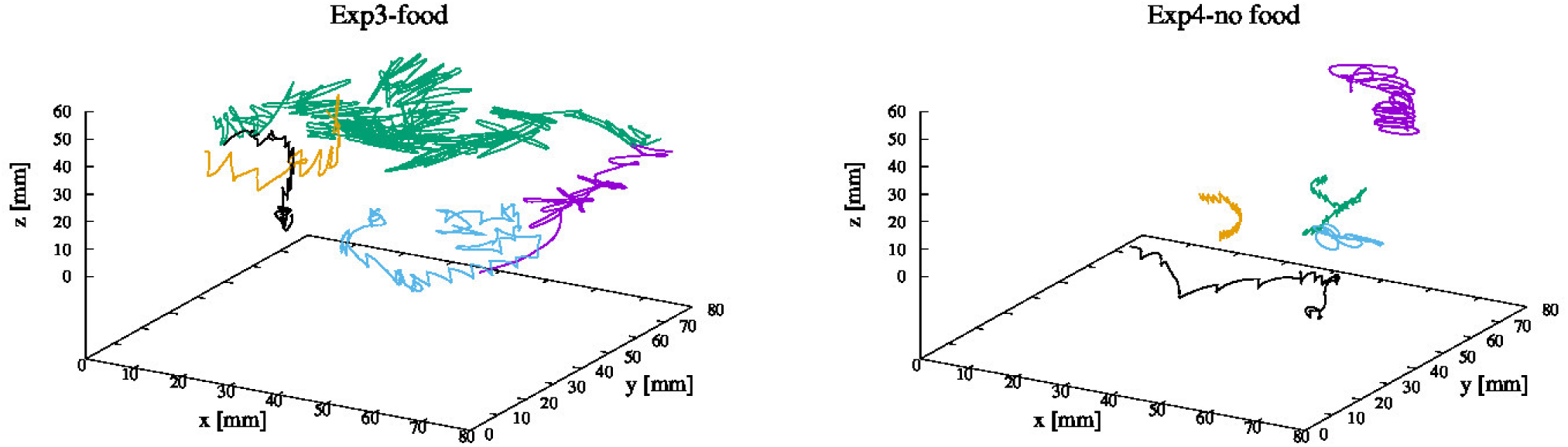
3D swimming trajectories of *Clausocalanus furcatus* in the presence (a) and absence (b) of food. Trajectories last as follows. Panel a: 24.4 *s* (violet), 241.6 *s* (green), 100.6 *s* (cyan), 54.8 *s* (orange), 60.4 *s* (black). Panel b: 32.6 *s* (violet), 61.6 *s* (green), 10.6 *s* (cyan), 56.2 *s* (orange), 64.6 *s* (black).

**Figure 2:**
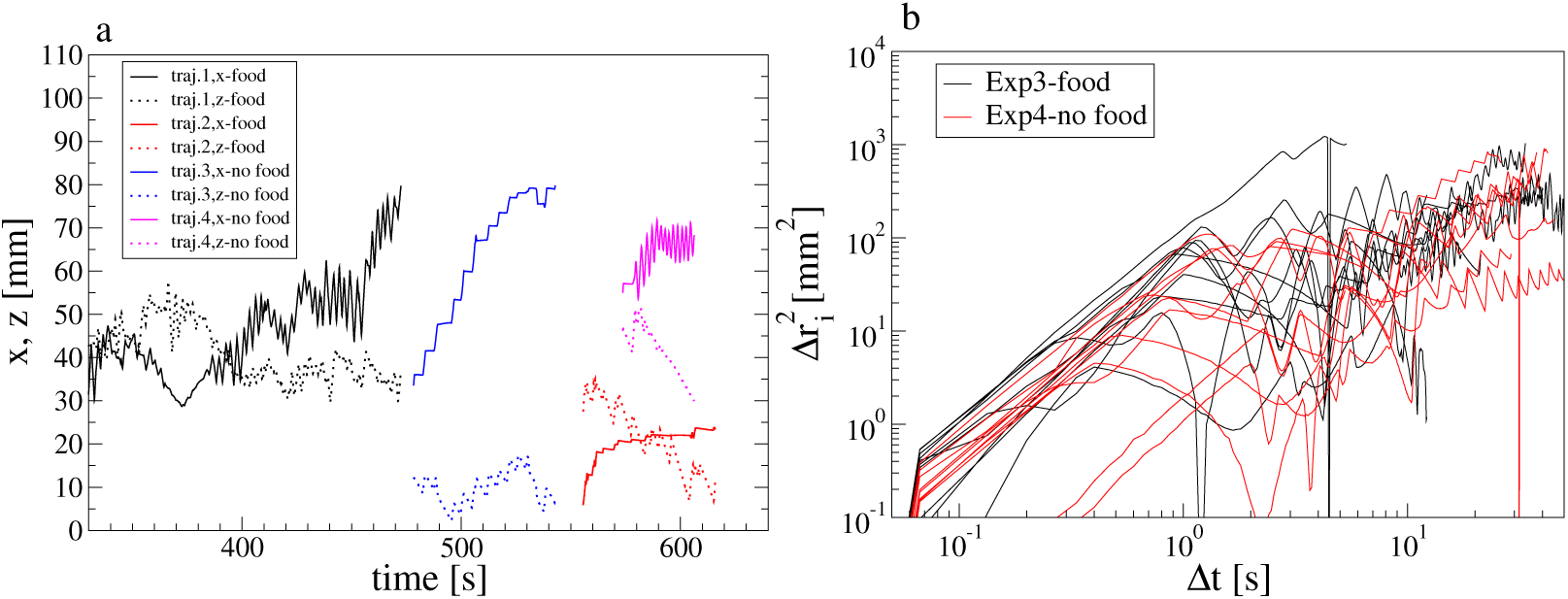
(a) Examples of spatial coordinates in the horizontal plane (x axis) and in the vertical direction (z axis). Different colors indicate different trajectories. For each trajectory, solid and dashed lines indicate the x and z component, respectively. (b) Examples of 3D square displacements, 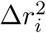, as a function of time and in the presence and absence of food, as indicated.

Figure 3 shows the lag-time dependence of the total, 3D MSD in the presence and absence of food. Some common properties emerge in all the investigated systems. At short time, Δ*t* ≤ 1 *s*, the MSD in creases ballistically, 〈Δ*r*^2^(Δ*t*)〉 = *D_b_*Δ*t^α^*, with *α* ≃ 2 and *D_b_* being an effective diffusion coefficient that characterizes the ballistic regime. This is consistent with the fact that *C. furcatus* moves mainly straight and at constant speed on short time scales ^10^^;^^6^^;^^23^^;^^26^. *D_b_* is generally smaller in the absence of food, suggesting the existence of a conservative mechanism: the organisms tend to reduce their speed and, therefore, the energy dissipation when food is scarce. *D_b_* is slightly larger in Exp3 (higher food concentration) than in Exp1, which suggests a smooth onset of such a conservative mechanism. However, the difference between Exp1 and Exp3 is too small to draw robust conclusions, whereby a larger number of experiments at different food concentrations should be required for understanding to what extent does the speed increases with food availability.

**Figure 3:**
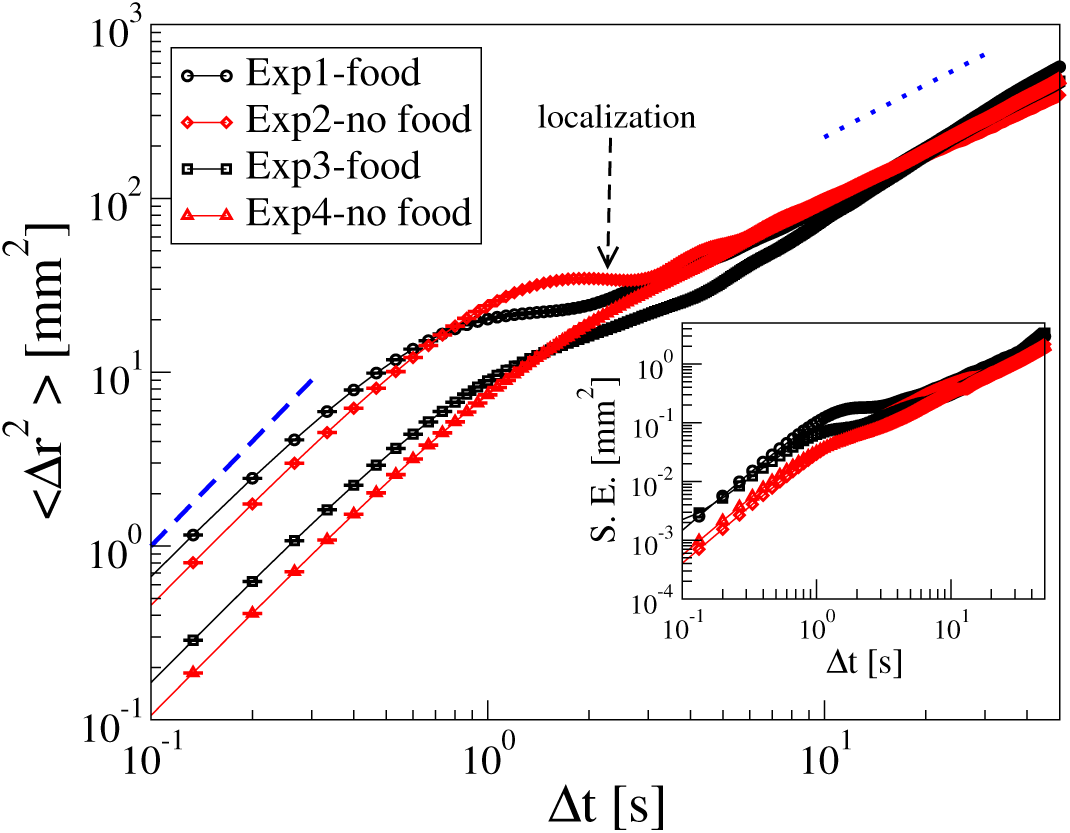
Three-dimensional mean square displacement of *Clausocalanus furcatus* as a function of the lag-time, 〈*Δr*^2^(Δ*t*)〉, for the four experiments in the presence and absence of food, as indicated. The standard error, *S.E.* is reported both as error bars and in the inset. The dashed and dotted lines are guide to the eye corresponding to a ballistic, 〈Δ*r*^2^(Δ*t*)〉 ∝ *t*^2^, and diffusive, 〈Δ*r*^2^(Δ*t*)〉 ∝ *t* behaviour, respectively.

Another common property emerges at longer time, Δ*t* ≥ 10 *s*, and consists in the approach to a diffusive regime, 〈Δ*r*^2^(Δ*t*)〉 = *D*Δ*t*^γ^, with *γ* ≃ 1. Accordingly, on large time and length scales, *C. furcatus* moves similarly to a random walker. The typical trajectory duration is too short for fully entering the diffusive regime and robustly estimate the diffusion coefficient *D* but the data overlapping at long time suggests that *D* is poorly affected by the presence of food. While MSDs with and without food appear similar at short and long time scales, distinctive features become manifest when focusing on intermediate time scales, Δ*t* ∊ [1*s*; 10*s*]. Indeed, for the two experiments in the presence of food, the ballistic and the diffusive regimes are separated by a clear localized regime, with the MSD attaining a temporary plateau, 〈Δ*r*^2^(Δ*t*)〉 ≃ *const* or equivalently, 〈Δ*r*^2^(Δ*t*)〉 ∝ Δ*t^β^*, with *β* ≃ 0. We estimated the root MSD at the plateau to be *λ_p_* = 5.8 *mm* and *λ_p_* = 4.5 *mm* for Exp1 and Exp3. Conversely, a smooth crossover from ballistic to diffusive regime is observed when food is not present, *i.e.* the exponent *β* gradually decreases from 2 to 1. The intermediate time difference emerging in the presence and absence of food is well beyond the statistical uncertainty, as clarified by the standard error on the MSD data, 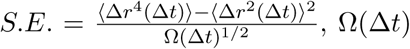 being the size of the statistical ensemble, reported both as error bars and, separately, in the inset.

Inspection of single organism trajectories suggested that swinging motion may appear more frequently when focusing on the horizontal projection of the trajectories, rather than along the vertical direction^6^. To clarify this point, Fig.4 a and b show the vertical and the horizontal contribution to the mean square displacement, 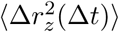 and 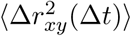, respectively. Regardless of the presence of food, the intermediate time plateau is definitely suppressed along the vertical direction. As a consequence, the localization observed in the 3D MSDs must arise from the horizontal contribution only. Indeed, Fig.4 b confirms that the intermediate time plateau appears even more clearly in 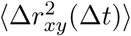. It is worth noticing that *C. furcatus* suspended in water do swim in a non-isotropic environment. While the horizontal component of the interaction is hydrodynamic only, a gravitational contribution is present along the vertical direction, since these organisms are not density matching, but slightly heavier that water (1.0281 – 1.0316*g cm*^−3 27^). Accordingly, it is likely that the optimal strategy to explore different directions may not be the same.

**Figure 4:**
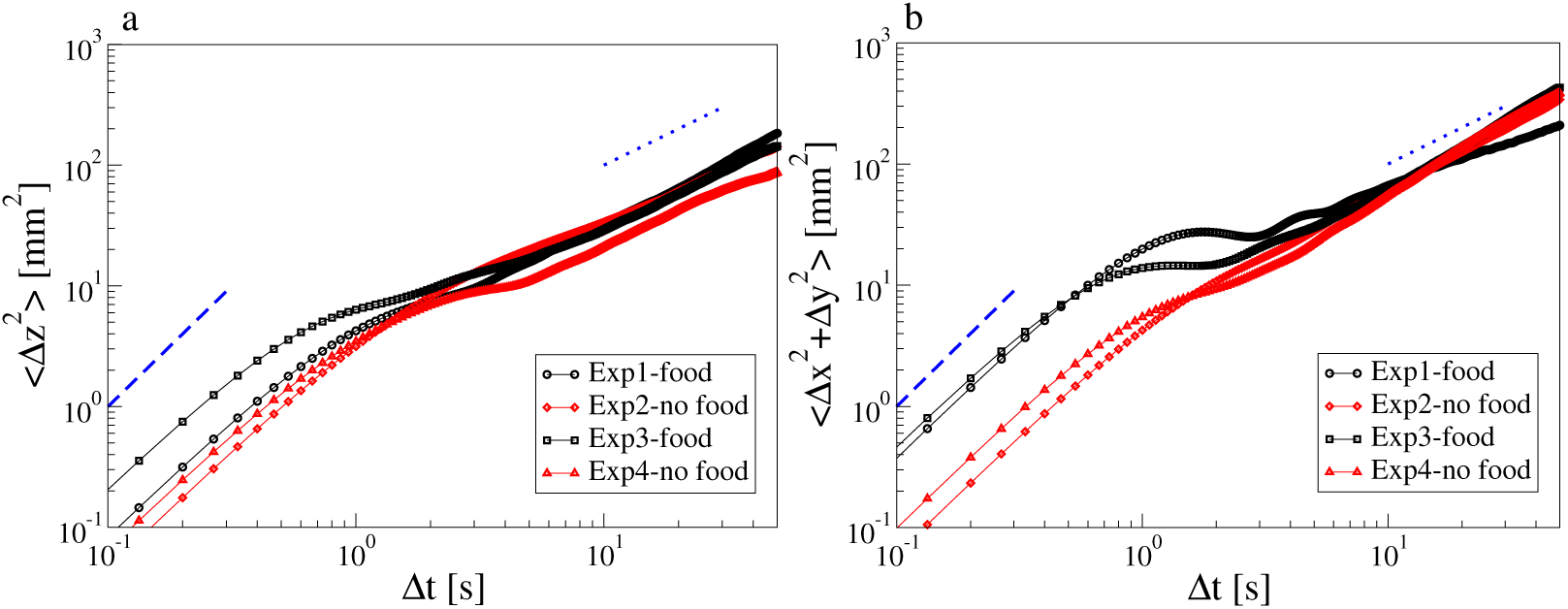
For the four different experiments, mean square displacement (a) along the vertical (*z*) direction and (b) in the horizontal (*x, y*) plane. The dashed and dotted lines are guide to the eye corresponding to a ballistic, 〈Δ*r*^2^(Δ*t*)〉 ∝ *t*^2^, and diffusive, 〈Δ*r*^2^(Δ*t*)〉 ∝ *t* behaviour, respectively.

To further characterize the dynamics of the system, we investigated the dynamic correlation function *F*(*λ* Δ*t*), commonly used to characterize the relaxation dynamics of liquids and soft materials and to spot out the presence of localized dynamics^25^^;^^28^^;^^29^. In particular, we focused on the horizontal plane, where the difference between experimental conditions was the most marked.

Figure 5 shows *F*(*λ* Δ*t*) as function of Δ*t* and at different values of the probe length in the range *λ* ∊ [1; 35]. In the absence of food (panel b), *F*(*λ* Δ*t*) shows a gradual decay that, as expected, becomes slower for increasing probe lengths and eventually vanishes over timescales exceeding the experimental time, as observed at the largest *λ* values. The overall scenario in the presence of food is similar (panel a), but in a range of probe lengths, *F*(*λ* Δ*t*) shows a two step decay with an intermediate time plateau. This is a typical signature of temporary localization, since it indicates that displacements of length *λ* are typically anti-correlated in the form of back and forward movements, over timescales of the order of the plateau duration. A similar signature is, indeed, observed in glassy liquids and colloidal suspensions^30^^;^^31^^;^^32^^;^^33^. We do observe this plateau in a range of timescales and probe lengths, which is fully compatible with the plateau of 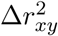 (see 4). Much larger (or much smaller) probe lengths are expected to be insensitive to localization. This is exactly what is observed at the largest investigated values of *λ* where the plateau is, indeed, not present. Overall, this result confirms that the presence of a temporary localized regime in the horizontal plane represent a clear-cut benchmark for predicting whether food is present or not in a given environment.

**Figure 5:**
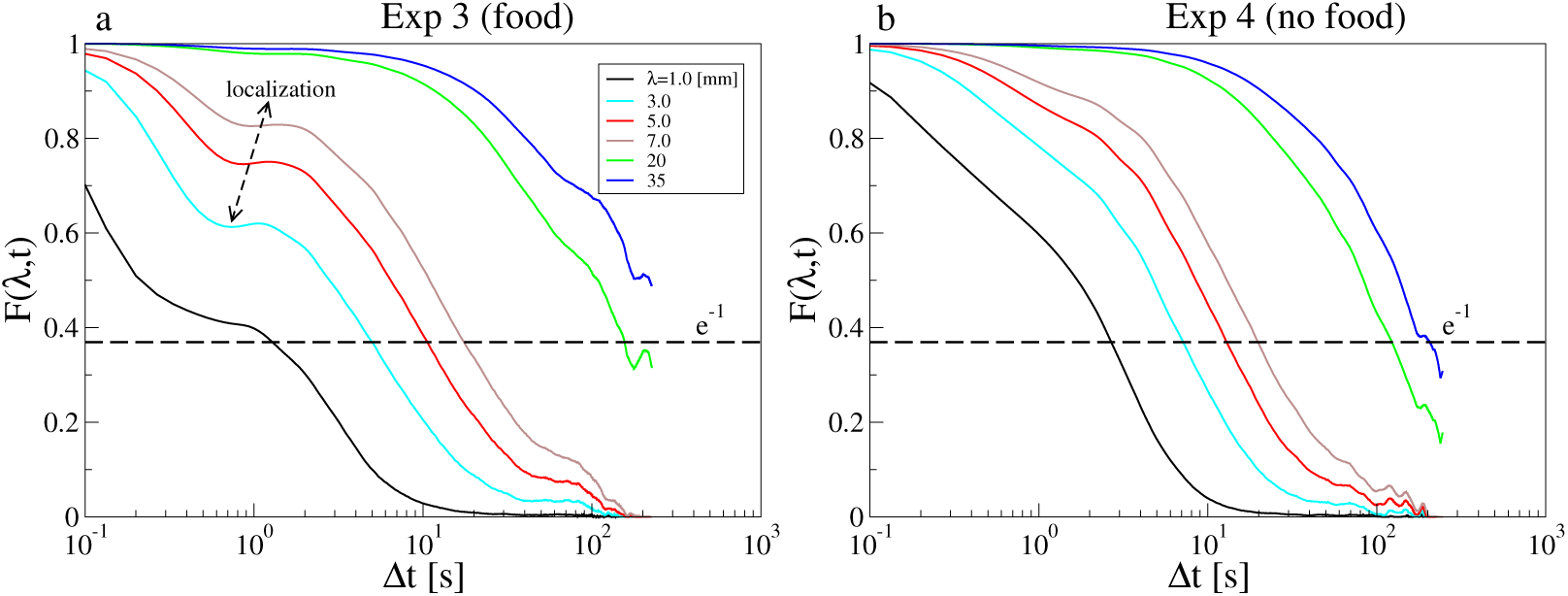
Lag-time dependence of the dynamic correlation function relative to the horizontal plane, *F*(*λ* Δ*t*), at several values of the probe length, *λ* as indicated. Panels (a) and (b) refer to two experiments, in the presence and absence of food, respectively.

From the decay of *F*(*λ* Δ*t*), we measured the relaxation time, *τ_λ_*. Figure 6 shows a plot of *τ_λ_* as a function of *λ* for two experiments in the presence and absence of food. In both cases, *C. furcatus* trajectories were compatible with a ballistic and a diffusive behaviour in the limit of short and long probe lengths respectively, therefore confirming the results from MSD. It is worth noticing that, although both 〈Δ*r*^2^(Δ*t*)〉 and *τ*(*λ*) provide a spatio-temporal description of the motion, the resulting information are not exactly the same. Indeed, MSD is controlled by the faster particles, while the relaxation time by the slowest ones. Thus, major differences may arise, especially in systems characterized by an heterogeneous distribution of mobility.

**Figure 6:**
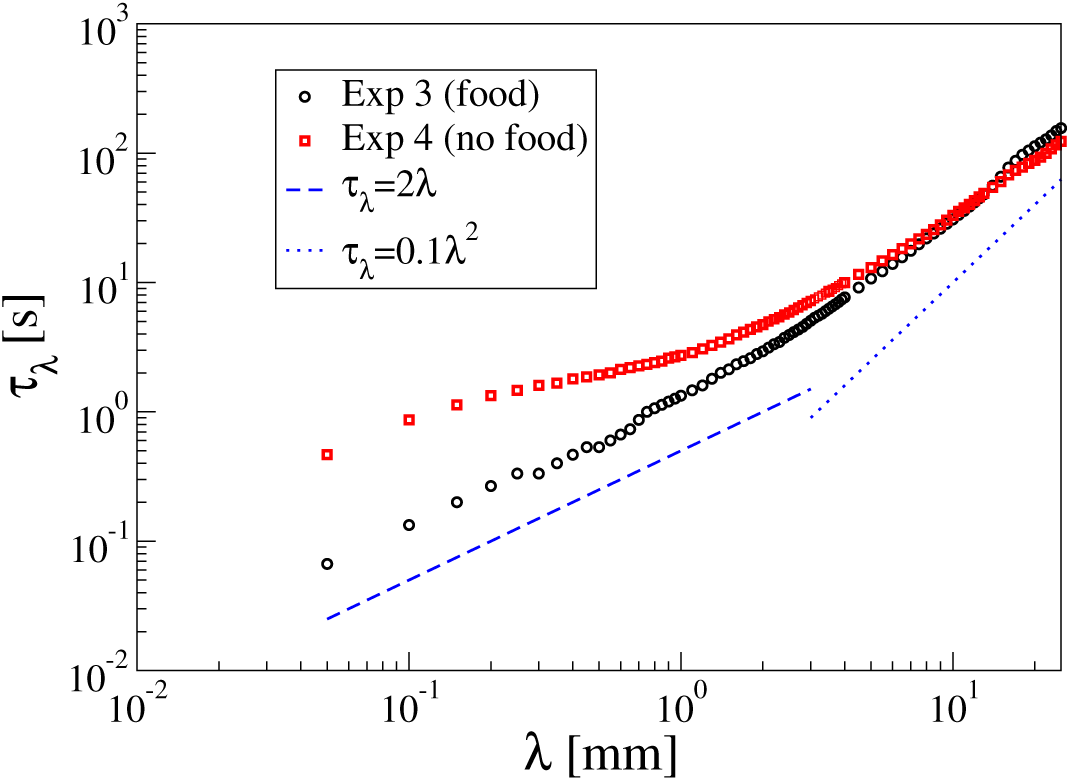
Relaxation time *τ_λ_ vs.* the probe length *λ* for two experiments, in the presence and absence of food, as indicated. The dashed and dotted line are guides to the eye corresponding to ballistic, *τ_λ_* ∝ *λ* and diffusive regime, *τ_λ_* ∝ *λ*^2^, respectively.

## 4 Discussion and Conclusions

Living organisms can enhance the encounter rates with prey (or mates) by moving along patterns with specific statistical properties^34^, although this behaviour is counterbalanced by the demand of reducing the hazardous meeting with predators. Planktonic organisms, like many other active systems ^13^^;^^14^^;^^15^^;^^16^, obtain such a tradeoff combining motions with ballistic and diffusive features^12^^;^^35^. Ballistic behaviour has been suggested to be an optimal strategy for small-scale processes, especially provided that the characteristic length of interaction with prey is smaller than the correlation length, *i.e.* the distance travelled before correlations in the velocity direction vanish. Conversely, diffusive behaviour emerges at large scale, when the correlation length is much shorter than the predator perceptive distance^12^.

Previous works on the small calanoid copepod *Clausocalanus furcatus* revealed a variety of swimming patterns and highlighted the unique characteristics of its behaviour with respect to other copepods^10^^;^^23^. Incidentally, it is worth noticing that these works are based on lateral view videos and, therefore, on the the 2D trajectory projections in the vertical plane. Some novel and major kinematic features in the motion of *C. furcatus* are found to be mainly manifested in the horizontal plane, and, therefore, are discernible only using specifically conceived 3D observation setups. For example, such an experimental approach allowed to identify semi-quantitatively recursive swimming patterns, including fast looping connected by sharp turns, and up and sink patterns combined with circular patterns in the horizontal plane^6^^;^^26^.

The analyses performed in our work deepen our knowledge of the behavioural mechanisms by which *C. furcatus* adapts to its environment, suggesting high degree of plasticity with respect to the external conditions. The MSD and the dynamic correlation function support the scenario of a ballistic motion at small spatio-temporal scales and diffusive regime at large scales and unveil novel benchmarks of the swimmer motion. First, we do find that, in a food rich environment, the transition between these regimes is characterized by a temporary localization, whereas the ballistic to diffusive crossover is smooth when food is absent. In addition, this localized regime is clearly identified in the horizontal plane, while it is almost indiscernible in the vertical direction.

The localization separating the ballistic and the diffusive regimes in the presence of food indicates that *C. furcatus* explores intensively a small region for several seconds and then randomly moves to a new neighboring region. While the mechanism behind subdiffusion and localization are, in general, manifold^36^, similar temporary localized motion is commonly observed in inert systems, such as glassy liquids and colloidal suspensions, as well as in experiments and simulations of many active, biological systems, including epithelial cell tissues, micro-swimmers and ant colonies^37^^;^^38^^;^^39^^;^^40^. While localization in such examples is strictly related to crowding and accompanied by large dynamical correlation (*e.g.* swarming), this is not the case for the system considered here, and the similarity is to be understood in a purely phenomenological sense. Indeed, in our case, the concentration of organisms is low, the encounter rates negligible, and, accordingly, crowding does not play a relevant role. Conversely, collisions of *C. furcatus* with food items and consequent stop for eating them may be a simple and interesting hypothesis for the observed localization. In this case, the root MSD at the plateau, *λ_p_* should be determined by the average distance, *l_f_* between food items, that can be simply estimated by the food concentration, 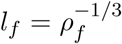, assuming a constant and homegeneous distribution of food ^1^. Using this approximation, we find *l_f_* = 1.26 *mm* and *l_f_* = 0.58 *mm* for Exp1 and Exp3, respectively. By comparison with *λ_p_* we can conclude that the prediction is qualitatively in agreement with the trend of the two experiments, but significantly smaller than the measured values of *λ_p_* i.e. *λ_p_/l_f_* ≃ 4.6 and *λ_p_/l_f_* ≃ 7.6 for Exp1 and Exp3, respectively. This suggests that such a simple mechanism may be relevant for the diffusion process but, overall, further and more complex behavioral processes are probably at work. In addition, inspection of single trajectories (see Fig.2) suggests that localization is mainly due to forward and backward movements, which are difficulty explained only in terms of collision with prey.

We speculate that in the presence of food, *C. furcatus* explores a limited region by continuous looping (mostly in the horizontal plane)^6^, reducing path crossovers at the small scale to increase the encounter with prey^26^. By contrast, the diffusive behaviour at the large scale associates with high crossovers^26^, likely ensuring reduced encounters with predators. The reduction of *D_b_* with the concentration of food may be considered as a metabolic energy-saving mechanism that, coupled with the swimming dependent encounter rate with prey^41^, supports the successful development of C. furcatus populations in oligotrophic conditions^18^. In addition, the plateau in the MSD (localized regime) parallels the transition in the number of crossovers occurring at scales typical of the shift from prey/mates to predator encounter^26^. The volume of the localization region may be related to the concentration of food through a conservative mechanism, such as 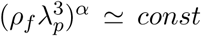. The simplest case, *α* = 1, qualitatively captures the trend of *λ_p_* in the two experiments with food, but is not quantitatively compatible with the results.

In other species observed so far in laboratory conditions, the transition between ballistic and diffusive motion is smooth^12^^;^^42^. The presence of the intermediate time plateau in *C. furcatus* is much likely associated to its typical motion behaviour, which can be considered unique among planktonic copepods.

Compared to direct inspection of each single trajectory, our approach allows to highlight quantitatively the emergent and dominant features of the motion, but also leads to an unavoidable loss of information. As a perspective, we plan to bridge average description and direct inspection of single particles by investigating other statistical descriptors that explicitly keep trace of the individual variability. Future perspectives include bridging average description and direct inspection of single particles by investigating other statistical descriptors that explicitly keep trace of the individual variability (*e.g.* Van Hove and waiting time distributions). In addition, a larger number of experiments at different food concentrations would elucidate the dependence of *D_b_* and *λ_p_* on *ρ_f_.*

## 5 Appendix

In this section, we provide a description of the averaging procedure adopted in this work. Our averaging approach is commonly used with particle tracking data from experiments on colloidal suspensions and other soft materials. Let us consider a generic quantity, *f_i_*(*t*_0_, *t*_0_ + Δ*t*), relative to the *i^th^* trajectory and depending on an arbitrarily chosen time origin, *t*_0_, and a time-lag, Δ*t* = *t* − *t*_0_. For example, 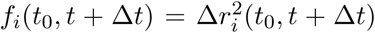. In this paper, 〈*f*(Δ*t*)〉 indicates that the average is performed over the ensemble of all trajectories and time-origins, *t*_0_, available in a given experiment. Precisely,
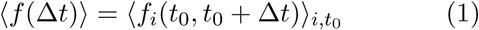

It is worth noticing that as Δ*t* increases, the size of the statistical ensemble, Ω(Δ*t*), decreases. Indeed, both the number of trajectories longer than Δ*t* and the number of available time-origins in each of these trajectories decrease, since the relation, *t*_0_ + Δ*t* ≤ *t*_*i*_, must hold for each trajectories of duration *t_i_*. Accordingly, average data become less meaningful and robust at large time, especially for Δ*t* > 50 *s*.

## 6 Acknowledgements

R.P. thanks the COST ACTION MP1305 *Flowing Matter* for inspiring this work. G.B. was supported by the Centre for Animal Movement Research (Can-Move) financed by a Linnaeus grant (no. 349-2007-8690) from the Swedish Research Council and Lund University

## Footnotes

1 A source of potential mismatch is due to the fact that our estimation is based on the initial value of the food concentration. As time passes and individuals eat more, food progressively decreases and, therefore, *l_f_* increases. Accordingly, we expect it to be a lower boundary for the real value of *l_f_.*

